# Perceptual insensitivity to higher-order statistical moments of coherent random dot motion

**DOI:** 10.1101/261370

**Authors:** Michael L. Waskom, Janeen Asfour, Roozbeh Kiani

## Abstract

When the visual system analyses distributed patterns of sensory inputs, what features of those distributions does it use? It has been previously demonstrated that higher-order statistical moments of luminance distributions influence perception of static surfaces and textures. Here, we tested whether the brain also represents higher-order moments of dynamic stimuli. We constructed random dot kinematograms where dots moved according to probability distributions that selectively differed in terms of their mean, variance, skewness, or kurtosis. When viewing these stimuli, human observers were sensitive to the mean direction of coherent motion and to the variance of the individual dot displacement angles, but they were insensitive to skewness and kurtosis. Observer behavior accorded with a model of directional motion energy, suggesting that information about higher-order moments is discarded early in the visual processing hierarchy. These results demonstrate that use of higher-order moments is not a general property of visual perception.

## INTRODUCTION

Perception emerges from the statistical analysis of sensory information (Helmholtz, 1867; Jazayeri & Movshon, 2006; Pouget, Dayan, & Zemel, 2000). What statistical features are used in this analysis? When processing the distribution of luminance across a static surface, human vision is sensitive to both lower- and higher-order moments. We perceive mean luminance as brightness, luminance variance over space as spatial contrast, and positively skewed luminance as “gloss” (Motoyoshi, Nishida, Sharan, & Adelson, 2007). It is largely unknown, however, whether higher-order moments also convey meaningful information in other visual domains.

One domain that lends itself well to experimentally measuring the influence of higher-order moments is visual motion. When viewing a collection of elements that move independently according to samples from a probability distribution, as in a random dot kinematogram, the visual system can extract a percept of coherent motion. Random dot kinematograms that are generated from a uniform or normal distribution appear to move coherently in the direction of that distribution’s mean (Watamaniuk, Sekuler, & Williams, 1989; Williams & Sekuler, 1984). Here, we ask whether the higher-order moments of the motion distribution correspond to other percepts. Based on prior studies of static textures (Kingdom, Hayes, & Field, 2001; Motoyoshi et al., 2007; Okazawa, Tajima, & Komatsu, 2015; Portilla & Simoncelli, 2000), it might be expected that they would.

We presented observers with a large random dot kinematogram and asked them to identify the location of an “odd” patch embedded within it. In the odd patch, dots moved with statistics that systematically and selectively differed from those in the rest of the stimulus. We found that human observers could detect odd motion only when either the mean or variance of the motion distribution was manipulated; in contrast, they were insensitive to changes in skewness and kurtosis. These results indicate that representation of higher-order moments is not a general property of the visual system.

## METHODS

### Observers and experimental setup

5 human observers (1 female; ages 19–36) participated in the experiment. Observers had normal or corrected-to-normal vision. One of the observers was author JA; the others were naive to the purposes of the experiment. Before participating, observers provided informed written consent; all experimental procedures were approved by the Institutional Review Board at New York University.

Observers were seated in an adjustable chair in a semi-dark room with chin and forehead supported before a cathode ray tube display monitor (21 inch EIZO; 75 Hz refresh rate; 1600 × 1200 screen resolution; 53.6 cm viewing distance). Viewing was binocular. Stimulus presentation was controlled with the Psychophysics Toolbox (Brainard, 1997) in MATLAB (The Mathworks). Gaze position was monitored at 1 kHz using a high-speed infrared camera (Eyelink; SR-Research).

### Odd patch detection task

Figure 1A shows a diagram of the behavioral task sequence. The observer initiated each trial by looking at a central fixation point (0.3° diameter). Following a variable delay (0.4 – 1 s, truncated exponential), a random dot kinematogram appeared within a 10° wide aperture surrounding the fixation point. Dots were white squares (93.1 cd/m^2^; 3 pixel × 3 pixel; 0.072° × 0.072°) drawn on a black background (0.89 cd/m^2^) with an average dot density of 100.2 dots per square degree per second. These parameters are similar to those used in other psychophysical and neurophysiological experiments (Britten, Shadlen, Newsome, & Movshon, 1993; Watamaniuk et al., 1989).

**Figure 1:**
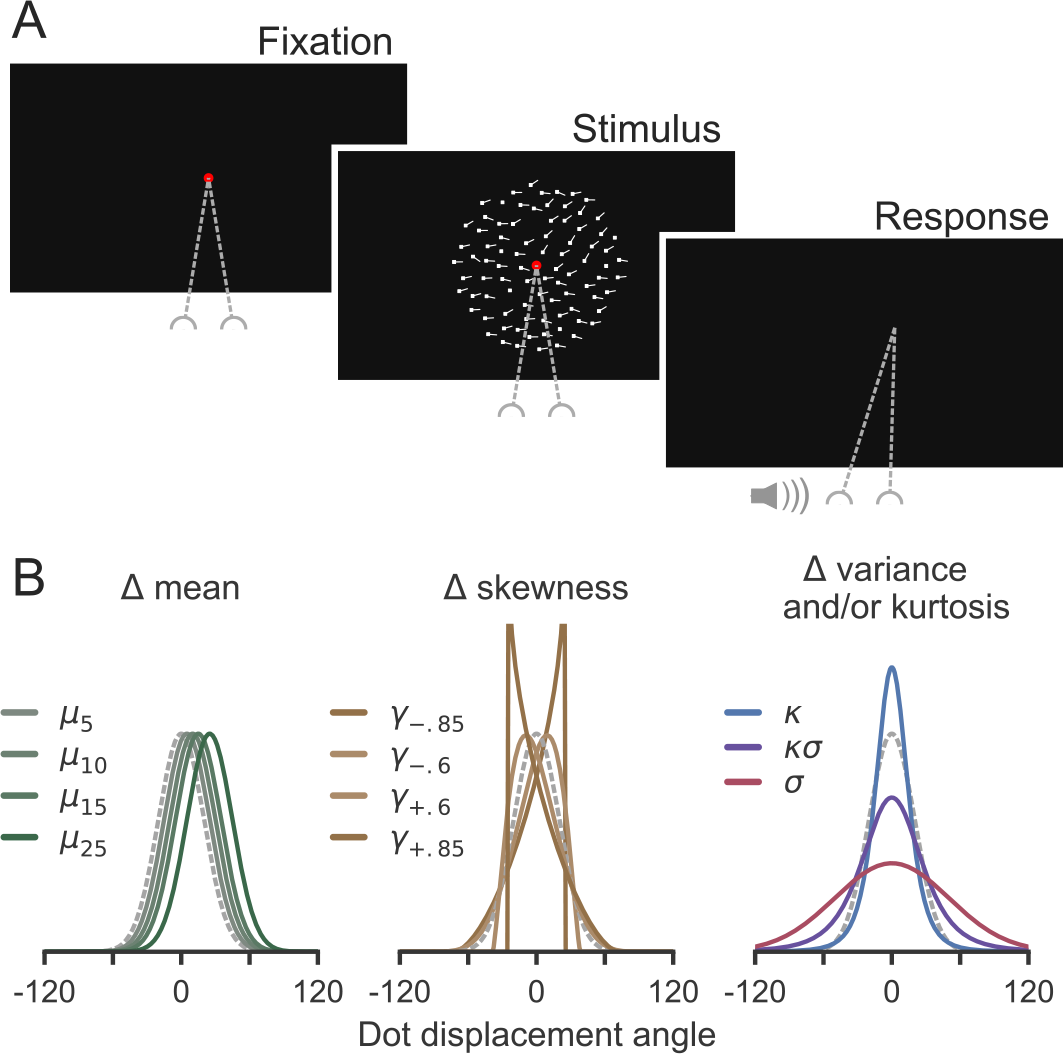
Experimental design. (A) Observers fixated while viewing a field of moving dots until they detected the presence of a patch with “odd” motion compared to the background. They indicated their detection when it occurred by making a saccade into the patch and received auditory feedback about their accuracy. (B) Generating distributions for dot displacements in different experimental conditions. Background motion was generated from a normal distribution and is shown on each plot with dashed gray lines.

On each screen refresh, dots were displaced coherently with probability 0.9 to create apparent motion at 5 degree per second. The remaining dots were redrawn randomly within the aperture. Random redrawing limited the half life of each dot to 6.5 frames (88 ms), preventing long streaks. The direction of motion for each of the coherently moving dots varied from one frame to the next. For the majority of dots (the “background”), the direction of displacement on a given frame was drawn from a Gaussian distribution (mean=0°, s.d.=20°). Dot motion within a specific circular aperture (the “odd patch”) was generated from one of several different distributions. The odd patch aperture (3° diameter) was always located in the top half of the display, and it was always centered at 3.5° eccentricity from the fixation point. Its radial position varied between 0° and 180° from trial to trial. To reduce adaptation, we added 180° to the mean of both the background and odd distributions on a random half of the trials.

Dot motion within the odd patch was generated using distributions that differed from background motion in terms of their mean, variance, skewness, and/or kurtosis while holding the other statistics constant (Figure 1B and Supplemental Movies). To accomplish this, we used the Pearson distribution system as implemented in the MATLAB function pearsrnd (Johnson, Kotz, & Balakrishnan, 1995). We parameterized the Pearson system to produce a diverse set of distributional shapes. Specifically, on ~19% of trials, the mean was varied to take values of 5°, 10°, 15°, or 25° (*μ*_5_, *μ*_10_, *μ*_15_, and *μ*_25_ conditions, respectively). On ~19% of trials, the skewness was varied to be either positive or negative across two magnitudes (±0.6 and ±0.85; *γ*_−.85_, *γ*_−.6,_ *γ*_+.6,_ and *γ*_+.85_ conditions). On ~19% of trials, the kurtosis was increased to 1802 accompanied by a small increase in standard deviation to 20.4°. (*κ* condition), and, on ~19% of trials, the same change in kurtosis was accompanied by a larger change (38°) in standard deviation (*κσ* condition). On ~19% of trials, the standard deviation alone was increased to 50° (*σ* condition). Finally, in ~4% of trials, the dot motion in the odd patch aperture was generated with the same statistics as the background (⌀ condition).

The observers were instructed to identify the location of the odd patch and to make a saccade to its location as soon as it was detected. Observers were required to maintain fixation without blinking until making their response. They were not provided any specific instructions as to what features should be used to identify the odd patch, but they were told that it would always be in the upper half of the display. The stimulus was turned off when gaze left a window around the fixation point (3° diameter), and the trial otherwise terminated if 10 s elapsed without a response (1.1% of trials). Feedback for correct and incorrect choices was provided through two distinct auditory tones. Observers were extensively trained on the task until their accuracy on the *μ*_&_ trials exceeded 80%; they were unaware of the specific criterion. Training was performed over multiple sessions and before beginning collection of the data reported here.

### Calculating sample statistics of dot displacement

There are two potential reasons that we could have failed to generate motion stimuli with distinct higher-order moments. First, dot displacements were generated using continuous coordinates in degrees, but to show the dots, those positions had to be discretized onto the grid of monitor pixels. Second, in any given trial, the observer sees a finite number of dot displacements, and so the ability to estimate the parameters of the motion distributions, particularly for the higher-order moments, might be limited by insufficient data. To evaluate these potential limitations, we must also consider that the Pearson system is defined on the real number line. Therefore, to determine how far our stimuli diverged from the theoretical values of the different distributions, it was necessary to evaluate the circular statistics of samples derived from those distributions. Sample circular statistics (Fisher, 1995) were computed using the CircStat and pycircstat libraries (Berens, 2009).

We first estimated limiting values for the generating moments by drawing large samples (N = 100,000,000) using each set of Pearson distribution parameters and computing the circular mean, standard deviation, skewness, and kurtosis. Next, to evaluate the possibility of recovering moment values from the dot stimuli themselves, we computed circular statistics from the stimuli shown on each trial. That is, we measured the angle of each coherent dot displacement after discretization into pixel coordinates and then calculated the sample statistics of that distribution. This computation excluded the final 300 ms of the stimulus, corresponding to the approximate non-decision time (i.e. the approximate sum of visual latency and saccade latency; Palmer, Huk, & Shadlen, 2005; Resulaj, Kiani, Wolpert, & Shadlen, 2009). We did this so that the statistics would correspond to the dot motion that was most likely used to make the decision on each trial.

Figure 2 shows sample statistics for each condition. It is clear that sample means derived from the stimulus closely tracked the generating distributions. The agreement between the generating and stimulus standard deviations was also close, although the latter slightly overestimated the former across all conditions. On conditions where we manipulated skewness or kurtosis, the sample statistics slightly underestimated the generating moments but were clearly distinct from both the background statistics and the odd patch statistics in other conditions.

**Figure 2:**
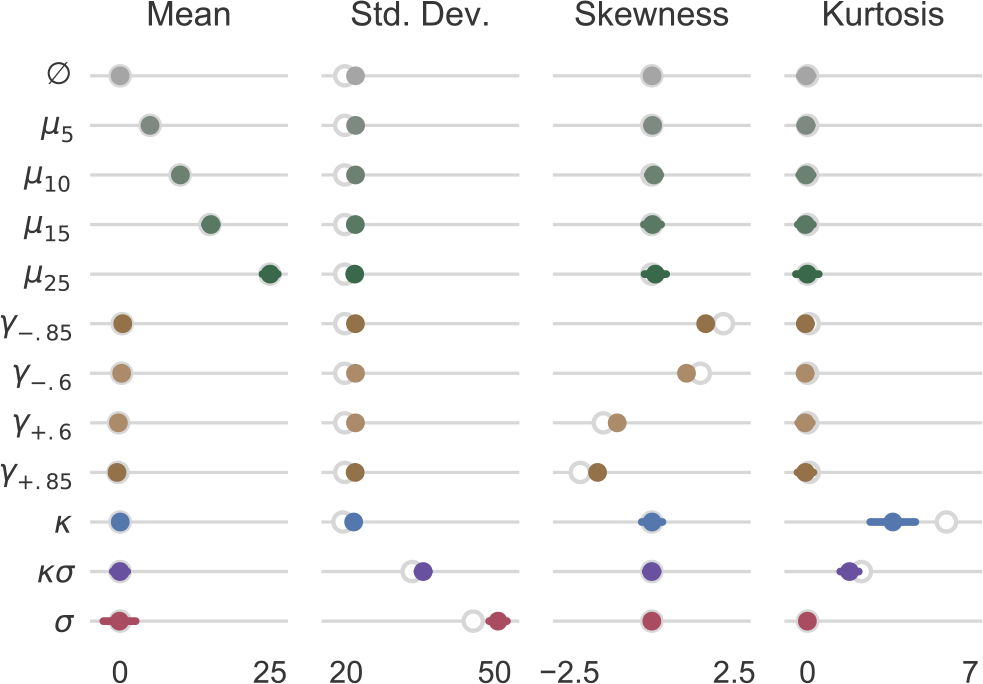
Sample statistics of dot displacement distributions. Open circles show the corresponding circular moment of the generating distribution at the limit of very large samples. Solid points and error bars show the means and standard deviations of the observed stimulus statistics across trials. Single-trial moments were computed using circular statistics based on the finite sample in each trial. Sample means and standard deviations are shown in degrees.

### Statistical analyses of behavior

Statistical analyses were performed on all trials with a valid response, which was defined as the eye position leaving the fixation window and landing between 1.75° and 5° from the fixation point. In total, the analyses involved 5349 trials. We used general linear models to evaluate the influence of the odd patch distribution statistics on both choice accuracy (i.e., the probability of making a saccade into the odd patch) and reaction time (RT). Specifically, we used logistic regression to analyze accuracy and linear regression to analyze RT. For these models, we used either the condition-wise means of the sample statistics computed on each trial as described above or, in separate models, the trial-wise statistics. Condition-wise and trial-wise models were otherwise structured the same way.

For choice accuracy, we characterized the effect of the displacement statistics with the function

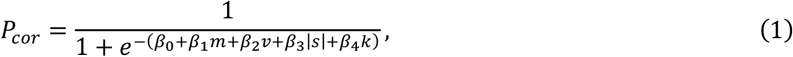

where *β*_I_ are regression coefficients and *m*, *v*, *s*, and *k* are the circular mean, standard deviation, skewness, and kurtosis. We used absolute skewness because we expected that only the magnitude, and not the direction, of skewness should be relevant. For each moment, the null hypothesis was that changes in the corresponding sample statistic would have no influence on the probability of detecting the odd patch (H_0_: *β_i_* = 0 for *i* > 0).

For RT, we characterized the effect of the statistics with the function

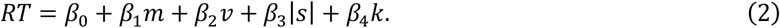

As above, the null hypothesis for each moment was that changes in the corresponding sample statistic would have no influence on the time taken to respond (H_0_: *β_i_* = 0 for *i* > 0).

Bar plots of behavioral results have error bars that show 95% bootstrap confidence intervals (Cumming & Finch, 2005). Confidence intervals were computed within each condition by resampling trials with replacement and estimating the statistic of interest across 100,000 iterations. The error bars correspond to the 2.5 and 97.5 percentiles of the resulting distribution. Plots of observer accuracy include an estimate of chance performance. This estimate is the ratio of the odd patch area to the area where a valid response could be registered (the upper half of the dot display from the outer bound of the fixation window to the outer bound of the dot aperture; see above).

### Motion energy model

We used arrays of oriented spatiotemporal filters to calculate the profile of directional motion energy produced by each stimulus distribution. The array of filters spanned direction selectivities from −90° to +90° in steps of 5°. We computed opponent motion energy using a pair of filers for each direction: one that was selective for that direction and one that was selective for the opposite direction.

Each directional filter was implemented as the sum of two space-time separable filters (Adelson & Bergen, 1985; Kiani, Hanks, & Shadlen, 2008). The spatial filters were even and odd symmetric fourth-order Cauchy functions:

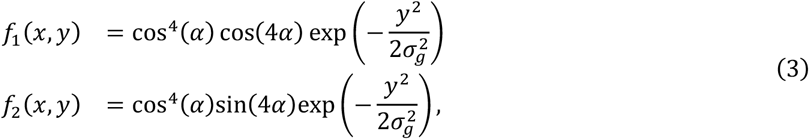

where *α* = tan^−1^(*x*/*σ*_c_). Along the *x* dimension, the filters resemble Gaussian-weighted cosine and sine functions. The envelope and period of the carrier functions are controlled by the order (fourth) and by *σ*_2_, which we set to 0.35°. They are windowed along the orthogonal (*y*) dimension using a Gaussian envelope with standard deviation *σ_g_* = 0.05°. The temporal filters were implemented as difference of Poisson functions:

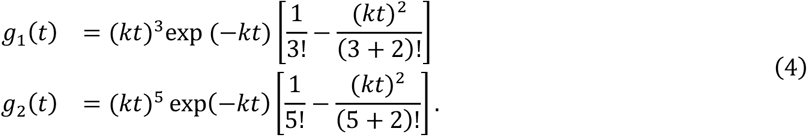

We set the time constant *k* of each temporal filter to 60. Together with the constants in the spatial functions, the filters implement a spatiotemporal frequency passband consistent with that measured in direction-selective middle temporal cortex (MT) neurons (Albright, 1984; Maunsell & Van Essen, 1983; Movshon, Newsome, Gizzi, & Levitt, 1988).

Direction selective motion filters were constructed by element-wise multiplication and summation of appropriate *f*_*i*_ *g*_*j*_ combinations. For example, *f*_1_ *g*_1_+ *f*_2_ *g*_2_ and *f*_2_ *g*_1_ − *f*_1_ *g*_2_pass motion in the +*x* direction (Eq. 3); this pair of filters is in space-time quadrature, which confers phase invariance.

To estimate motion energy, we first convolved each direction-selective spatiotemporal filter with a movie of the dot stimulus. Next, we squared and summed the quadrature pairs to produce a spatiotemporal image of local motion energy. These images were then summed over space, and net motion energy was computed by subtracting motion energy in the opposite direction. Finally, we took a mean over time within a window that excluded the latency of the motion energy filters (Eq. 4). This procedure was repeated across 500 different samples of the stimulus; each sample had a duration of 2 s (150 frames). The resulting motion energy values, averaged over samples, comprise a motion energy profile for each condition.

## RESULTS

### Perceptual sensitivity to motion statistics

Observers viewed a large random dot kinematogram and attempted to detect a small patch where the motion of the dots was “odd” compared to the background. The motion in the odd patch could differ from the background in terms of the mean, variance, skewness, or kurtosis of the dot displacement angles. Observers reported the odd patch by making a saccade to it as soon as it was detected. Figure 3A shows the probability of successfully detecting each kind of odd motion. It is immediately apparent that conditions where the odd patch differed from the background in terms of mean or variance produced detection rates above chance. In contrast, detection rates for odd patches that differed only in skewness or kurtosis were not different from chance. This pattern of behavior was consistent across individual observers (Figure 3B).

**Figure 3:**
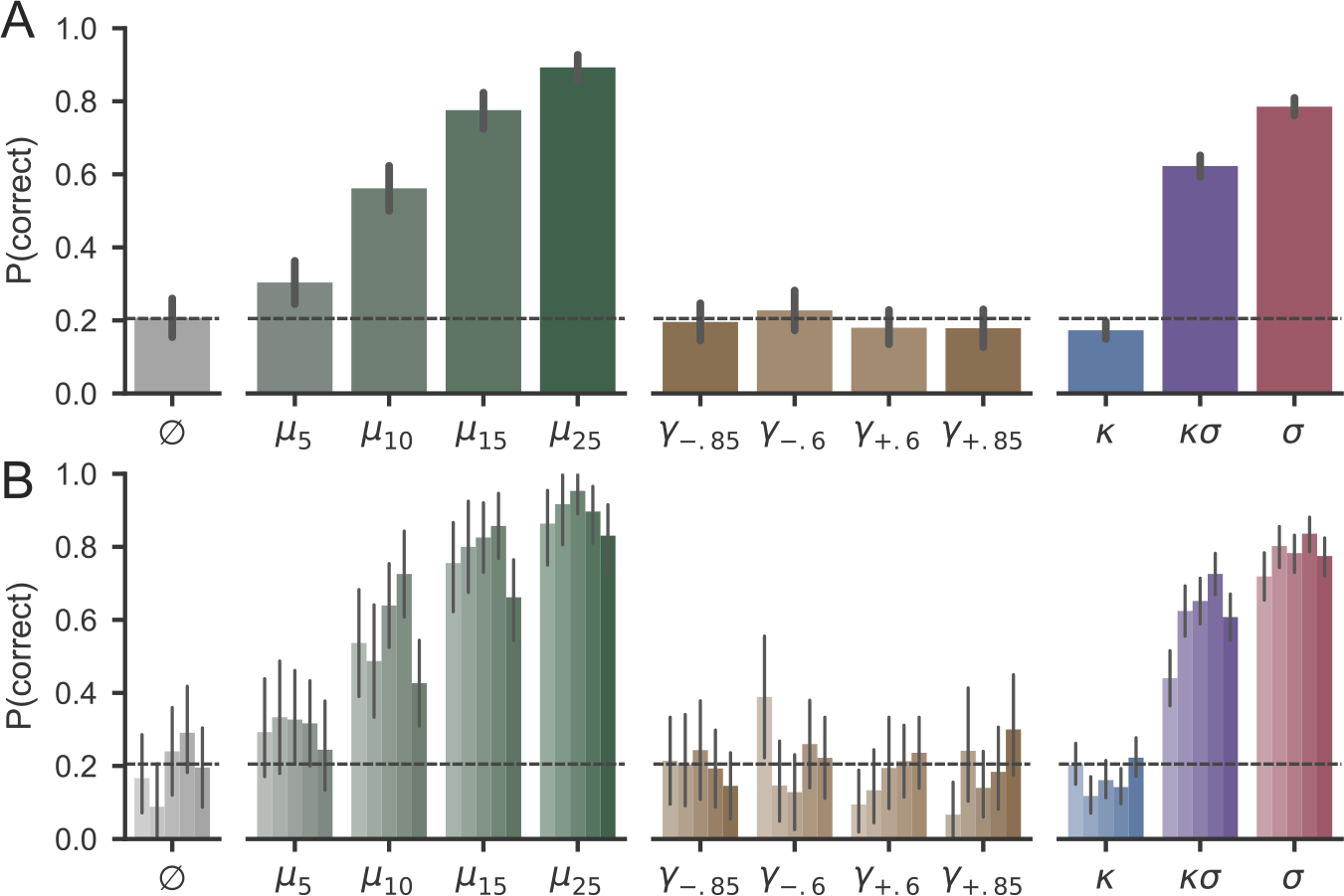
Probability of correctly detecting the odd patch in each condition. (A) Group results. Bar heights show mean accuracy across all trials; error bars show 95% bootstrap confidence intervals. The horizontal dashed line shows the expected value of chance performance (i.e., the probability that a random saccade would land within the odd patch; see Methods). (B) Individual observer results. Each bar shows a different observer with the same ordering across conditions. Conventions are otherwise as in panel (A).

We used logistic regression to evaluate these relationships quantitatively. The regression model (Eq. 1) confirmed that detection accuracy was significantly influenced by changes in mean (*β*_1_ = 0.15 ± 0.009, *z* = 16.63, *P* < 1e-8) and standard deviation (*β*_2_ = 0.10 ± 0.005, *z* = 22.35, *P* <1e-8) but not skewness (*β*_3_ = −0.073 ± 0.10 *z* = −0.73, *P* = 0.47) or kurtosis (*β*_4_ = 0.031 ± 0.035, *z* = 0.88, *P* = 0.38).

We wondered whether observers were sensitive only to odd motion with relatively large deviations from the background in terms of skewness or kurtosis. To evaluate this possibility, we fit two additional logistic regression models using trial-wise measures of the stimulus statistics, limiting each model to the conditions where either skewness or kurtosis were manipulated (Figure 4). In neither case did we find evidence for sensitivity to the higher moments: choice accuracy was not significantly influenced by trial-wise skewness on skew-manipulated trials (Eq. 1, *β*_3_ = −0.12 ± 0.27, *z* = −0.44, *P* = 0.66) or by trial-wise kurtosis on kurtosis-manipulated trials (Eq. 1, *β*_4_ = 0.10 ± 0.073, *z* = 1.40, *P* = 0.16).

**Figure 4:**
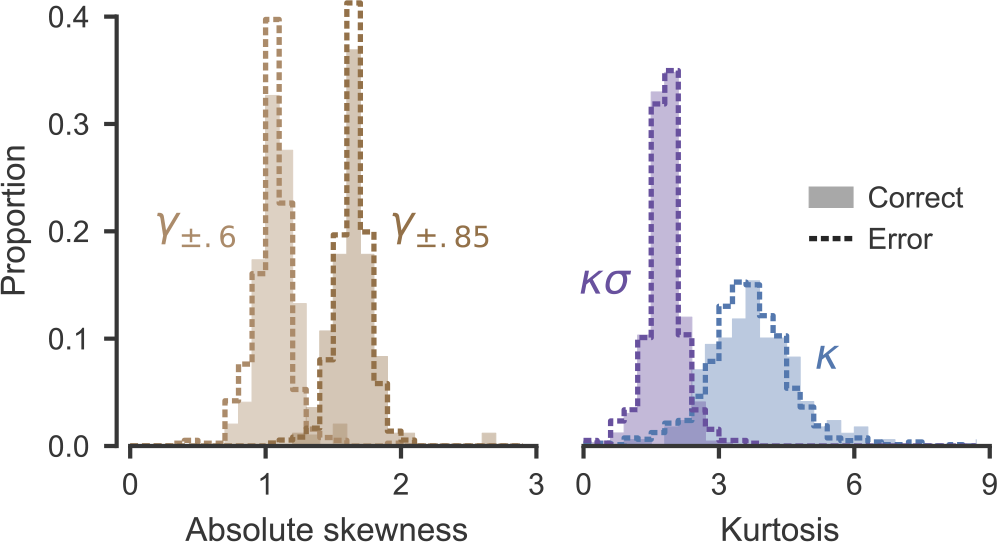
Trial-wise analysis of skewness and kurtosis. Histograms show the distribution of higher-order stimulus statistics depending on whether the observer produced a correct (shaded patch) or error (dashed line) response.

We also examined how motion statistics influenced the amount of time required to detect the odd patch. Figure 5A shows the mean RT for each condition. It is apparent that RTs were influenced by manipulations of mean and variance; in contrast, RTs on trials with a manipulation of skewness or kurtosis alone were similar to those on trials where the odd patch motion did not differ from the background (⌀ condition). A linear regression model (Eq. 2) confirmed these observations: RT was significantly influenced by mean (*β*_1_ = −0.11 ± 0.005, *t* = −21.25, *P* <1e-8) and standard deviation (*β*_2_ = −0.084 ±0.003, *t* = −26.40, *P* <1e-8) but not skewness (*β*_3_ = 0.054 ± 0.070, *t* = 0.77, *P* = 0.44) or kurtosis (*β*_4_ = −0.021 ± 0.025, *t* = −0.84, *P* = 0.40). This pattern of behavior was consistent across individual observers (Figure 5B).

**Figure 5:**
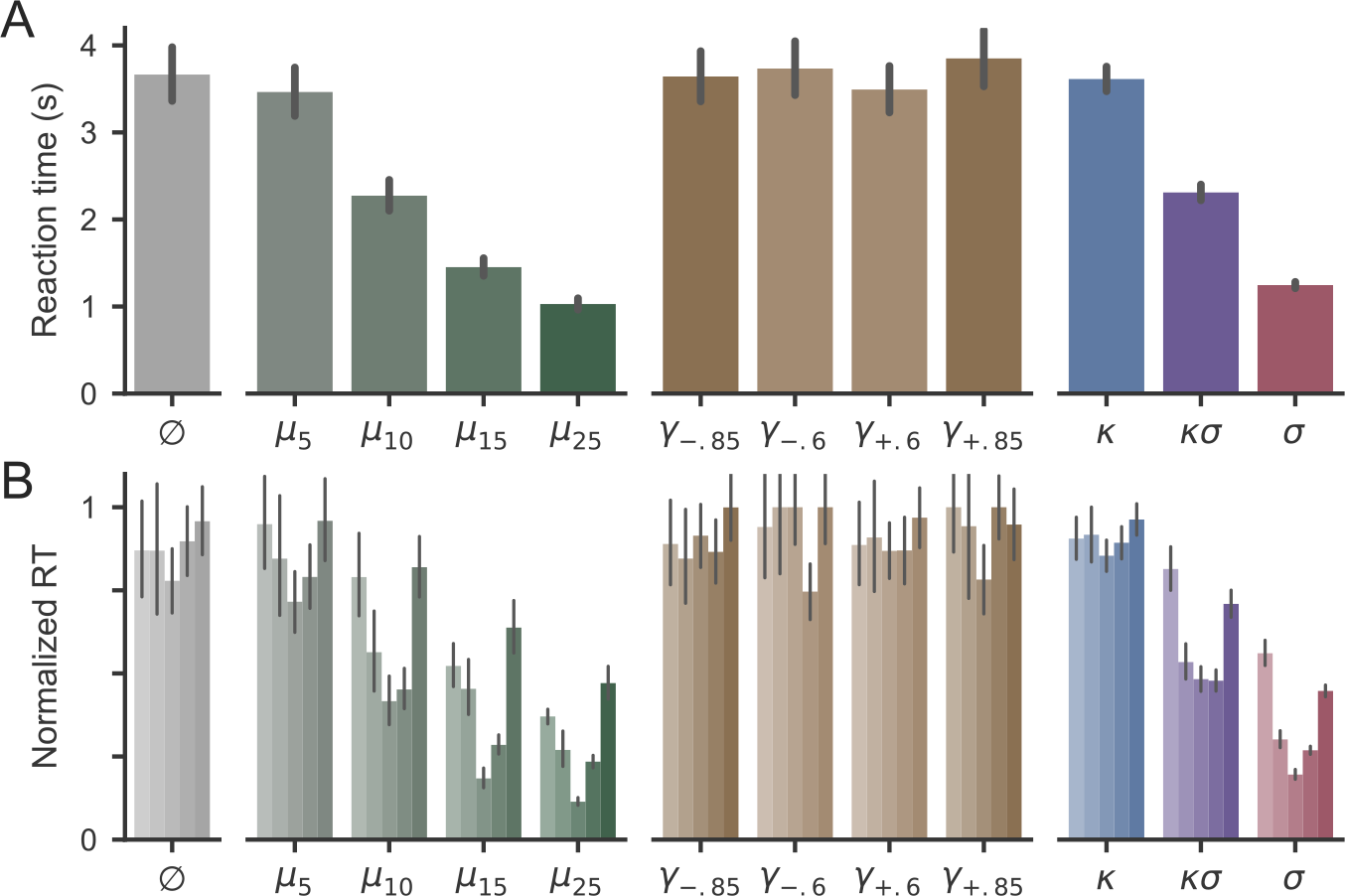
Reaction times in each condition. (A) Group results. Bar heights show mean RT across all trials; error bars show 95% bootstrap confidence intervals. (B) Individual observer results. Each bar shows a different observer with the same ordering across conditions. Each trial’s RT was normalized to the observer’s maximum condition-wise mean RT. Conventions are otherwise as in panel A.

### Motion energy model

Can a model of directional motion energy (Adelson & Bergen, 1985) account for these results? To evaluate this correspondence, we constructed an array of motion energy filters selective to different directions. The parameters for the filters were adjusted to match the direction tuning of motion selective neurons in MT cortex (see Methods). We compared the response profile of the motion energy filter array across the set of motion stimuli. These profiles are shown in Figure 6A. It is apparent that changes in the mean direction of coherent motion translate the motion energy profile and that changes in the variance of dot displacements broaden it. Consistent with the behavioral results, motion energy profiles for conditions where only skewness or kurtosis were manipulated do not appear appreciably different from the background motion energy profiles.

**Figure 6:**
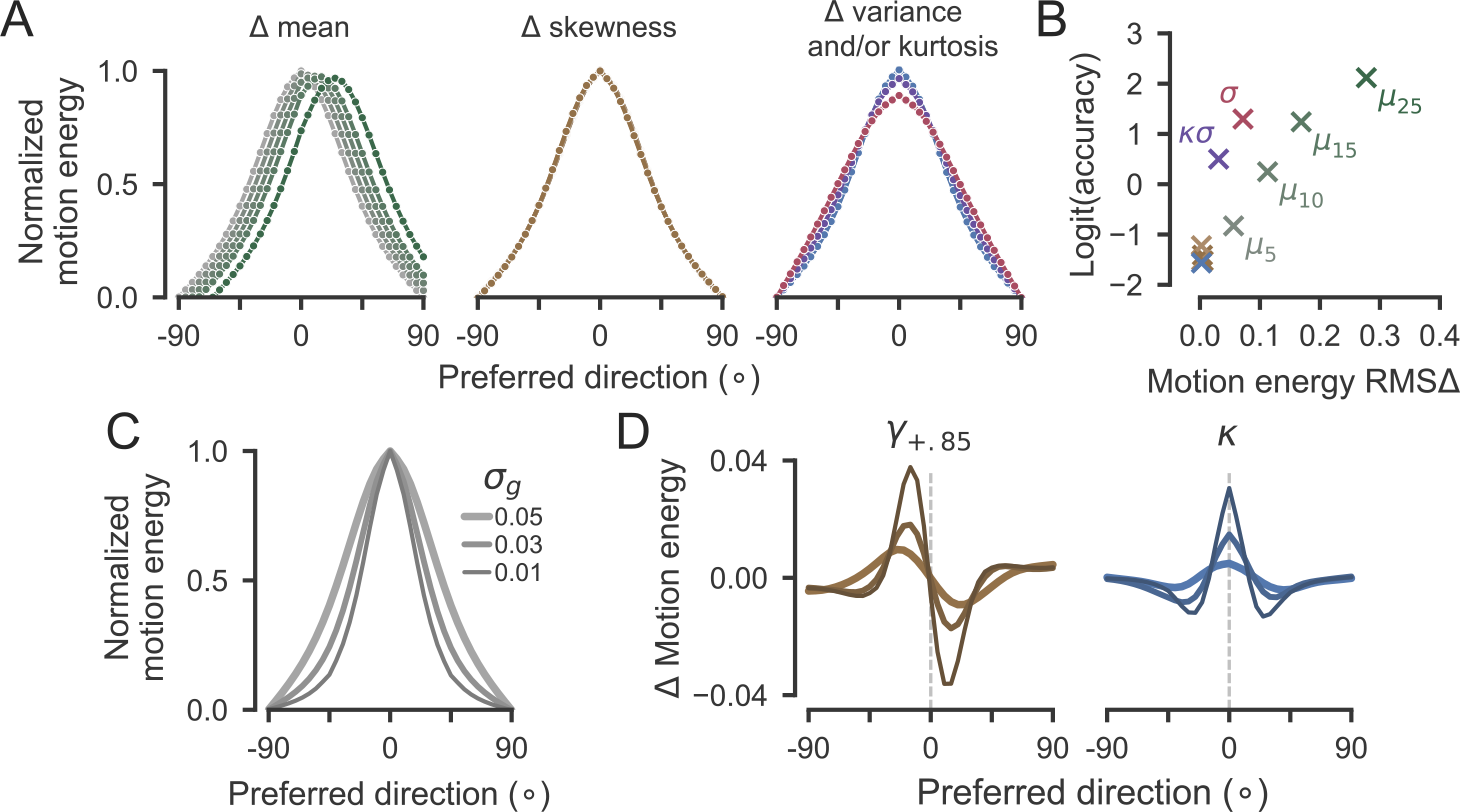
Motion energy analysis. (A) Normalized responses of motion energy filter arrays to each condition. All responses were normalized using the response of a filter with 0° direction preference to the background motion. The plots are organized as in Figure 1B. The response profile for background motion is shown in gray but is not visible on the skewness or kurtosis plots. (B) The relationship between detection accuracy and divergence between background and odd motion energy profiles across the filter arrays. Points corresponding to conditions with chance detection performance are unlabeled. Colors match the curves in panel A. (C) Direction tuning curves with three different tuning bandwidths. The curves in panel A were estimated using σ_g_ = 0.05 (Eq. 3). (D) Differences between normalized responses to high skew/kurtosis stimuli and to background motion in filter arrays with narrower direction tuning. Line widths correspond to the tuning curves in Panel C.

To relate the motion energy profiles to behavior, we computed the root-mean-square difference (RMSΔ) between each condition’s average motion energy profile and the motion energy profile of the background motion (Figure 6B). There are two observations to make about this plot. First, on mean-manipulated trials, the log-odds of detecting the odd patch increase roughly linearly with the divergence of the odd and background motion energy profiles. Second, relative performance on variance-manipulated trials is better than would be expected from the trend established by the mean-manipulated trials. While the RMS Δ measure is not intended as a formal decoding model, this relationship suggests that performance is supported by factors other than the shape of the time-averaged population response modeled by these motion energy filters, a point we return to in the discussion.

To determine whether the failure of the model to represent higher order moments could be attributed to its structure or to its parameterization, we constructed arrays of filters with narrower direction tuning curves (Figure 6C). We then applied these arrays to stimuli with high skewness and kurtosis. Narrowly-tuned filters were sensitive to higher-order moments, as evidenced by a shift in the peak corresponding to skew and a sharpening of the peak corresponding to kurtosis (Figure 6D). This result indicates that spatiotemporal filter arrays are not structurally insensitive to the higher-order moments of random dot motion. Nevertheless, when parameterized to match primate physiology, they are functionally insensitive to that information. Therefore, the inability to see differences of higher order moments offers insight into the objectives that the visual system has, and has not, evolved to satisfy.

## DISCUSSION

Perceptual systems are tasked with extracting meaningful information from patterns of sensory inputs. In naturalistic environments, these patterns may have complex distributions with features such as heavy or asymmetric tails. It has previously been shown that the perception of static surfaces uses some of this complexity to infer texture and material properties (Kingdom, Hayes, & Field, 2001; Motoyoshi et al., 2007; Okazawa, Tajima, & Komatsu, 2015; Portilla & Simoncelli, 2000). Here, we sought to determine whether higher-order moments are also used in the analysis of dynamic stimuli. Using well-controlled artificial stimuli, random dot kinematograms with different dot displacement distributions, we found that human observers were functionally blind to large changes in skewness or kurtosis when the mean and variance were identical to the background. This shows that, in at least some cases, the visual system discards useful sensory information to produce a compact internal representation that is limited to mean and variance. These results constrain theories of information processing in sensory systems.

Previous investigations of the relationship between dynamic stimulus statistics and perceptual experience have focused on higher-order correlations of spatiotemporal luminance patterns. These higher-order correlations arise in natural scenes, and modeling them is necessary to account for human motion perception (Hu & Victor, 2010; Nitzany & Victor, 2014). The present work differs from these previous efforts by considering the marginal statistics of individual elements that make up the random dot kinematogram. The perception of coherent motion in these stimuli is based on a representation of summary statistics (Watamaniuk et al., 1989), which is similar to the perception of static artificial textures (Victor, 1994), peripheral areas of natural scenes (Freeman & Simoncelli, 2011), and natural soundscapes (McDermott, Schemitsch, & Simoncelli, 2013). We found that humans were not sensitive to higher-order moments in artificial dynamic “textures”. Nevertheless, it is interesting to speculate about whether textures with such statistics exist in the natural world, perhaps in the motion defined by flocks of birds, herds of livestock, or crowds of humans.

By documenting a case where the visual system ignores higher-order moments, we have shown that the brain does not, as a rule, faithfully represent the distributional characteristics of its inputs. Nevertheless, there are domains in which higher-order moments are used for perception. Therefore, the task going forward is to determine when and why that is the case. Two alternate perspectives can guide such an investigation. From one, perceptual systems typically represent the higher moments of stimulus distributions, but in some cases, as with random dot motion, those statistics are discarded. Alternatively, it may be that stimulus distributions are typically reduced to a Gaussian representation and processed in terms of their mean and variance except in special cases where dedicated mechanisms for representing higher-order moments convey particularly useful additional information. Supporting the latter perspective, we note that Motoyoshi et al. (2007) propose a specific mechanism for the representation of luminance skewness, which may have developed because higher-order luminance statistics provide information about surface texture. Within the domain of dynamic information, it is important to note that we do not know whether our result will generalize to distributions over other aspects of translational stimuli, such as speed, or to other forms of motion, such as the dynamic textures that define optic flow.

Because we report a null result, it is natural to wonder whether larger changes in skewness or kurtosis might have been detected. While possible, we think this is unlikely because observers typically show high sensitivity for other aspects of the random dot stimulus. In our experiment, observers were above chance on trials with only a 5° change in mean direction, and direction discrimination thresholds in other paradigms can be as low as 2° (Watamaniuk et al., 1989). There is a long history of studying motion perception using random dot kinematograms, and their mean direction, speed, and coherence have close correspondence in neuronal responses (Britten et al., 1993). We cannot rule out the theoretical possibility that heavy-tailed motion could be detected in a stochastic stimulus created using a different parameterization or in a different class of spatiotemporal stimuli altogether. Yet even if such a stimulus could be created, it is striking that our observers were unable to detect large changes in skewness or kurtosis of the stimulus that we did use given their high sensitivity to changes in the first two moments. Therefore, our experiment represents a strong test, and refutation, of the hypothesis that the visual system always represents higher-order moments of sensory stimuli.

One challenge in using behavioral measurements to make inferences about information processing limitations is that it is often unclear at what stage potentially useful information may be lost. When behavior appears insensitive to some aspect of sensory input, it could be that sensory systems fail to preserve a representation of it or that inferential processes make poor use of that representation. In the present case, we are able to overcome this challenge by considering a formal model of motion perception (Adelson & Bergen, 1985). This model has close correspondence to the responses of direction-selective neurons in striate and extrastriate visual cortex (Albright, 1984; Maunsell & Van Essen, 1983; Movshon et al., 1988; Rust, Mante, Simoncelli, & Movshon, 2006). When using direction-selective spatiotemporal filters that matched the spatiotemporal tuning of the primate visual system, we found that the resulting motion energy profiles contained essentially no information about the skewness or kurtosis of the dot displacement distributions. This suggests that the visual system discards information about higher-order moments early in the processing hierarchy. Therefore, we can conclude that performance was limited at the sensory representation stage rather than by suboptimal inference. This sensory limitation arises from how inputs to direction-selective cells are combined to generate a representation of spatiotemporal energy.

More elaborate models of motion processing can account for other physiological phenomena, such as neural selectivity for pattern motion (Rust et al., 2006; Simoncelli & Heeger, 1998). These models build more complex representations from component elements that are equivalent to the motion energy profiles we estimated. Therefore, they should not be able to recover higher-order moments that are absent from the simpler representation. Nevertheless, considering models of later-stage motion processing may help to explain why performance on variance-manipulated trials exceeded what would have been expected from the motion energy profiles alone. One important factor to consider is that we examined the distribution of motion energy across an array of filters tuned to the speed of coherent motion in the background, yet increasing the variance of individual dot displacement angles will also decrease the average displacement in the mean direction. Therefore, a more complete mechanistic model of our task would likely need to represent the joint distribution of motion energy across a population of filters with varying direction and speed tuning.

In building a more complete model, it will also be necessary to consider how the observer represents and decodes momentary evidence to form a decision about the location of the odd patch. Our motion energy analysis focused on time-averaged differences in the motion energy profiles, but the observers were performing an RT task, and RTs were fastest in conditions with the largest divergence from the background motion. It is likely that the dependence of RT on the odd motion statistics can be attributed to multiple sources. Specifying a mechanistic model of odd motion detection will require answering several currently-open questions. First, does performance arise from a single strategy or from a mixture of condition-dependent strategies? Perhaps bottom-up recognition supports detection when odd motion differs strongly from the background, but top-down search must be engaged when the difference is more subtle. Second, are candidate odd patches subjected to an evidence accumulation process that integrates multiple samples of odd motion before committing to a decision? We are actively pursuing these questions to more fully understand the mechanisms that link stimulus statistics to behavior in the odd patch detection task.

If higher-order moments do exist in natural dynamic textures, the evolution of the visual system may have sacrificed the chance to see them because computing with compact representations of Gaussian statistics confers several benefits. An observer of motion is usually concerned with tracking the path of rigid bodies; indeed, assuming Gaussianity may help the visual system individuate multiple sources of motion that otherwise generate a platykurtic response in a population with broad direction tuning curves (Treue, Hol, & Rauber, 2000). More generally, Gaussian assumptions reduce processing demands for probabilistic computations because they require operations only on scalar representations of a distribution’s location and width. And because probabilistic computations on Gaussian inputs produce Gaussian outputs, this architecture can simplify inferential procedures. Nevertheless, a representation that is limited to lower-order moments poses a challenge to fully-Bayesian theories of neural computation, which would require more complete representations of probability distributions that include detailed information about their tails. Therefore, our results demonstrate a constraint that can inform future theories of how sensory information is encoded and decoded when using perception to guide behavior.

## ACKNOWLEDGEMENTS

We thank Chris Cueva and Bill Newsome for inspiring discussions and for their contribution to the conceptual development of this experiment in its early stages. We additionally thank Gouki Okazawa and Braden Purcell for comments on an earlier draft of the manuscript. This work was supported by a Whitehall Foundation Research Grant (2014-12-51), a Sloan Research Fellowship, and a McKnight Scholar Award to RK; MLW was supported by the Simons Foundation as a Junior Fellow in the Simons Society of Fellows (527794).

